# Leveraging foundation models to dissect the genetic basis of cluster compactness and yield in grapevine

**DOI:** 10.1101/2025.04.02.646891

**Authors:** Sadikshya Sharma, Jose Munoz, Efrain Torres-Lomas, Jerry Lin, Hollywood Banayad, Yaniv Lupo, Veronica Nunez, Ana Gaspar, Dario Cantu, Luis Diaz-Garcia

**Author notes:** Contributed equally (Sadikshya Sharma, Jose Munoz).

## Abstract

Grape cluster compactness is a key trait that influence fruit quality, yield, and disease susceptibility. Understanding the genetic basis of this trait is essential for optimizing vineyard management and improving grapevine cultivars. In this study, we performed quantitative trait locus (QTL) mapping to identify genomic regions associated with cluster architecture and yield components in a bi-parental population derived from *Vitis vinifera* cv. Riesling × Cabernet Sauvignon. A total of 138 full-sibling progeny were evaluated over two growing seasons at Oakville, Napa Valley, California. Traditional yield-related traits were measured, including cluster number, total cluster weight, and average cluster weight. Additionally, an image-based phenotyping pipeline leveraging the foundation model Segment Anything Model (SAM) was employed to segment individual berries, measure their size and shape, and compute cluster compactness with minimal manual intervention. Trait correlations revealed that compact clusters tended to have a higher berry count but smaller berry size, highlighting the role of compactness in modulating cluster structure. Heritability estimates varied across traits, with berry dimensions and compactness displaying moderate to high heritability, indicating strong genetic control. Two parental linkage maps were constructed using a pseudo-test cross strategy. QTL mapping identified multiple loci associated with cluster architecture and yield components, with several stable QTLs detected across both years. Notably, a QTL for cluster compactness was found in both seasons on chromosome 1 in Cabernet Sauvignon. Other stable QTLs were associated with berry size (chromosomes 6 and 17) and berry count (chromosome 5 in Cabernet Sauvignon and chromosome 7 in Riesling). Additional QTLs were detected in a single year, reflecting the influence of environmental variation. Our findings provide valuable insights into the application of foundation models requiring no prior training and minimal intervention for high-quality segmentation and enhance our understanding of the genetic architecture of cluster compactness and yield traits. The genomic regions identified in this study offer promising targets for breeding programs aimed at improving grape quality and disease resistance.

## Introduction

Grape cluster architecture and compactness are two interrelated traits essential for vineyard management and the genetic improvement of grapes, as they influence yield, fruit quality, and susceptibility to pests and diseases [1-4]. Cluster architecture determines how the berries are arranged within a cluster and the distribution of free space [2,5]. Cluster compactness, on the other hand, is defined as the ratio of the volume of cluster components—such as the berries and the rachis—to the overall volume of the cluster [2,5]. Understanding the implications of cluster compactness is crucial, as its physical configuration can significantly impact berry development and the microenvironment within the cluster. When clusters are compact, berries are in close contact with one another, which can alter their physical and physiological properties. For instance, tight contact can inhibit the proper development of the waxy cuticle, compromising its function as a protective barrier against pathogens [6-8]. Furthermore, the inner portion of compact clusters has restricted airflow, resulting in higher humidity and temperature, creating a more favorable environment for pathogens [3,7]. In addition to reduced airflow, there is also reduced solar radiation reaching the inner berries of compact clusters, which affects berry ripening and composition [7].

Several studies have sought to identify the genetic factors underlying cluster architecture and compactness using a range of phenotyping methods, including both qualitative and quantitative approaches. One commonly used qualitative method relies on a set of descriptors that categorize clusters based on visual characteristics [3]. The most widely adopted system is the set of descriptors developed by the International Organization of Vine and Wine (OIV), which includes standardized criteria for evaluating various traits of the grapevine, including cluster morphology. For example, OIV descriptor 204 assesses cluster density, classifying clusters into five categories ranging from very loose, where berries are clearly separated with many visible pedicels, to very dense, where berries are visibly deformed due to compression. Additional descriptors, such as OIV 208 (cluster shape) and OIV 209 (number of cluster wings), can be used in combination to characterize overall cluster architecture. While OIV descriptors offer a relatively fast and low-cost means of phenotyping, their accuracy is often limited by the evaluator’s subjectivity [3]. When multiple evaluators are involved, differences in interpretation can introduce inconsistency and increase error. In contrast, quantitative phenotyping is generally preferred for statistical and genetic analyses, as it provides more precise, reproducible data that better supports the dissection of complex traits [2,9].

The analysis of 2D and 3D images of grapevine clusters offers a powerful approach for measuring cluster architecture and compactness. Techniques in this field range from traditional image thresholding for object identification to advanced machine learning models, some specifically trained on grapevine data, others based on more generalizable foundational models with little or no crop-specific training. These image-based methods enable the extraction of high-dimensional phenotypic traits, enhance the accuracy of characterizing complex traits like cluster architecture, and offer excellent scalability for large datasets. Examples of cluster imaging and image analysis include the study by Cubero et al. (2015), which developed a technique for measuring cluster compactness that was highly consistent with the OIV descriptor 204 [10]. Their approach utilized semi-automated image segmentation to remove the background from the images and segment the clusters into berries and rachis. These segmented images were subsequently used to collect morphological measurements and create models for predicting cluster compactness. Similarly, Underhill et al. (2020) employed high-throughput imaging to quantify cluster compactness and identified key quantitative trait loci (QTLs) related to cluster structure [3]. More recently, Torres-Lomas et al. (2024) demonstrated the application of the Segment Anything Model (SAM) for grape cluster image segmentation, enabling automated object identification without additional training [2]. Their study applied SAM to over 3,500 cluster images, generating more than 150,000 berry masks with spatial coordinates, which allowed comprehensive cluster architecture and compactness analyses. These examples illustrate the effectiveness of image-based phenotyping in providing precise, reproducible data that can enhance genetic analysis and breeding strategies for grapevine traits.

Cluster architecture and compactness are complex traits shaped by a combination of environmental conditions, genetic factors, and vineyard management practices [7]. Numerous studies have aimed to uncover their genetic basis, identifying a wide array QTLs distributed across the grapevine genome. For example, Correa et al. (2014) identified several QTLs associated with rachis architecture, including a major locus for rachis length located near the microsatellite marker VMC2D9 on chromosome 9, using an F1 biparental population derived from a Ruby Seedless × Sultanina cross [11]. Similarly, Richter et al. (2019) mapped over two dozen QTLs related to traits such as pedicel length, rachis weight and length, peduncle length, berry volume and number, shoulder length, cluster weight, and OIV 204 score, in an F1 population from a GF.GA-47-42 × Villard Blanc cross [5]. In another study, Underhill et al. (2020) combined manual and image-based phenotyping to identify QTLs associated with cluster compactness, berry weight, and rachis length, revealing key loci across multiple chromosomes [3]. Together, these studies highlight the polygenic and highly complex genetic architecture underlying grapevine cluster traits, emphasizing the need for advanced phenotyping and mapping strategies to fully dissect their genetic control.

In this study, we performed QTL mapping using an F1 mapping population derived from a cross between two pure *Vitis vinifera* cultivars with contrasting cluster architectures: Riesling and Cabernet Sauvignon. Our goal was to identify genomic regions associated with yield components and cluster architecture by leveraging the Segment Anything Model, or SAM. Using SAM, we processed over 7,000 2D cluster images, generating nearly 330,000 individual berry masks, enabling precise measurements of berry morphology, spatial distribution, and cluster compactness, all without requiring additional model training. This methodology allowed us to successfully identify multiple QTLs linked to cluster architecture, compactness, and yield, with several QTLs consistently detected across two growing seasons. These findings offer valuable insights into the genetic determinants of cluster architecture and compactness, paving the way for breeding strategies aimed at enhancing fruit quality and reducing disease susceptibility.

## Results

### Digital phenotyping for cluster compactness

To characterize grape cluster architecture and compactness, we phenotyped 414 vines from 138 progenies of a Riesling × Cabernet Sauvignon cross over two consecutive growing seasons, 2023 and 2024. This mapping population was grown in Oakville, Napa Valley, California—a premier winegrowing region in the United States.

We collected two sets of phenotypic data: traditional yield-related traits, including the number of clusters per vine, total cluster weight per vine, and average cluster weight; and digitally phenotyped traits, including berry number, projected area, length, and width. From these, we calculated the compactness index, which is defined as the ratio of the sum of individual berry projected areas to the convex hull surrounding the entire cluster.

Across both seasons, a total of 7,483 cluster images were acquired, with each cluster imaged from two angles to maximize berry visibility. Using the Segment Anything Model along with a previously developed algorithm for berry segmentation [2], we generated 329,469 berry masks (an example of a processed cluster is shown in Figures 1A and 1B). While a single image underestimates the true berry count due to occlusion, as demonstrated in our previous work and this study, a linear regression-based correction factor was applied to improve the accuracy of berry number estimates (Figure 1C). To quantify compactness, we calculated the ratio between the sum of all berry masks (i.e., total projected berry area) and the convex hull, defined as the smallest convex polygon encompassing all berry masks (Figure 1D). To visually validate this approach, we ranked all clusters by compactness index and selected the 30 least compact (top) and 30 most compact (bottom) clusters for comparison (Figure 1E). The gradient from less to more compact clusters is evident, with loosely arranged berries in clusters at the top of the panel and densely packed berries in those at the bottom.

**Figure 1.**
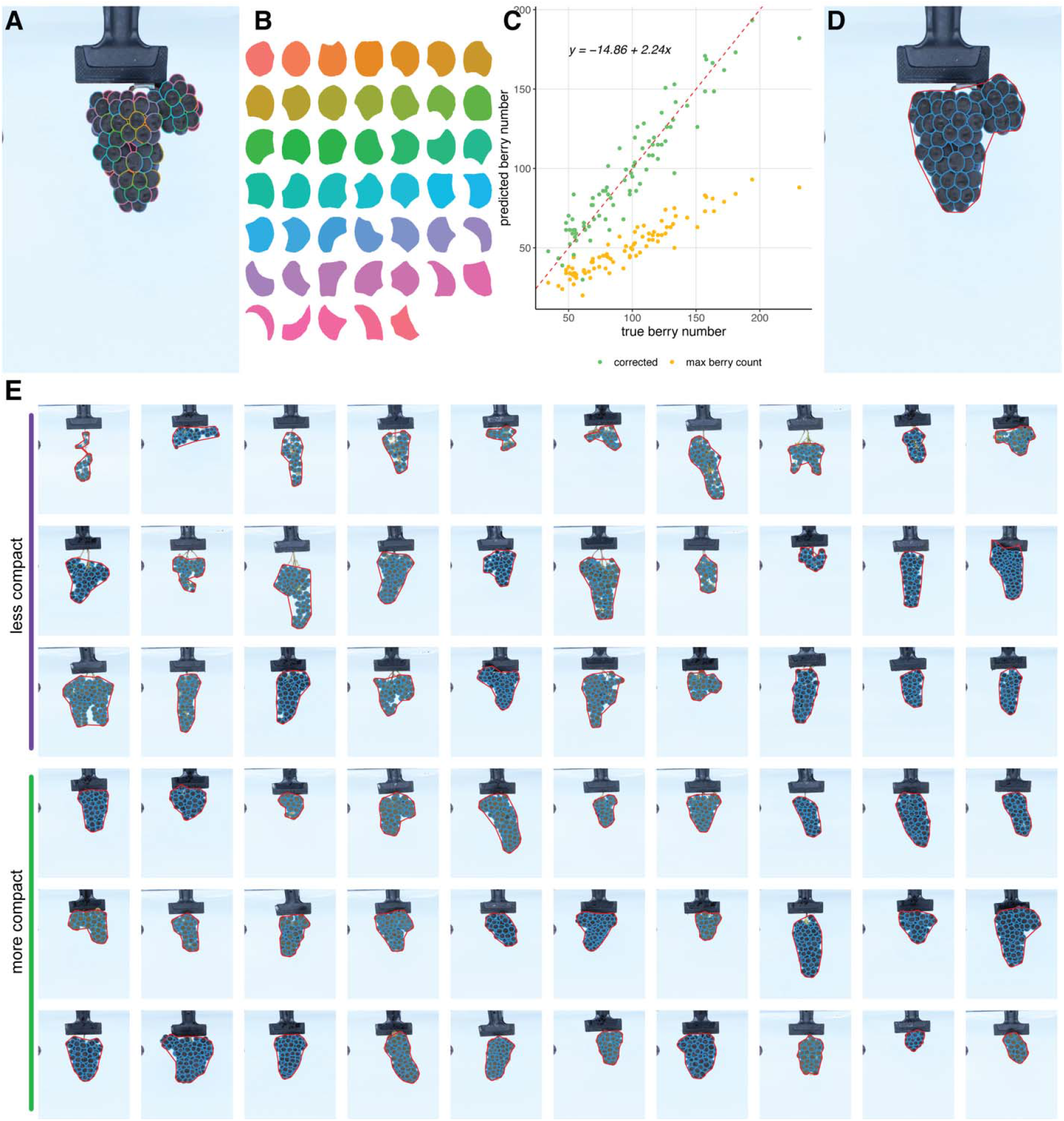
Identification of berry segments using the Segment Anything Model (SAM). **(A)** Example of individual berry identification from a grape cluster. **(B)** Berry masks generated from the cluster image in panel A. **(C)** Correlation between actual and predicted berry counts from SAM, using 84 clusters. Green points represent corrected counts, adjusted using a linear model based on the angle with the highest berry count. **(D)** Cluster compactness is calculated as the ratio of the total berry mask area to the area of the convex hull (outlined in red). **(E)** Examples of clusters with low and high compactness indices.

### Variation in yield and cluster traits across years

Both yield components and image-derived (digital) traits displayed continuous distributions with strong year-to-year consistency (*R*^*2*^ = 0.43 to 0.65; Figure 2A). Furthermore, transgressive segregation was observed across all traits. The two parental cultivars displayed expected phenotypic patterns: Riesling produced smaller, more compact clusters with fewer and smaller berries, while Cabernet Sauvignon produced larger, looser clusters with more and larger berries.

**Figure 2.**
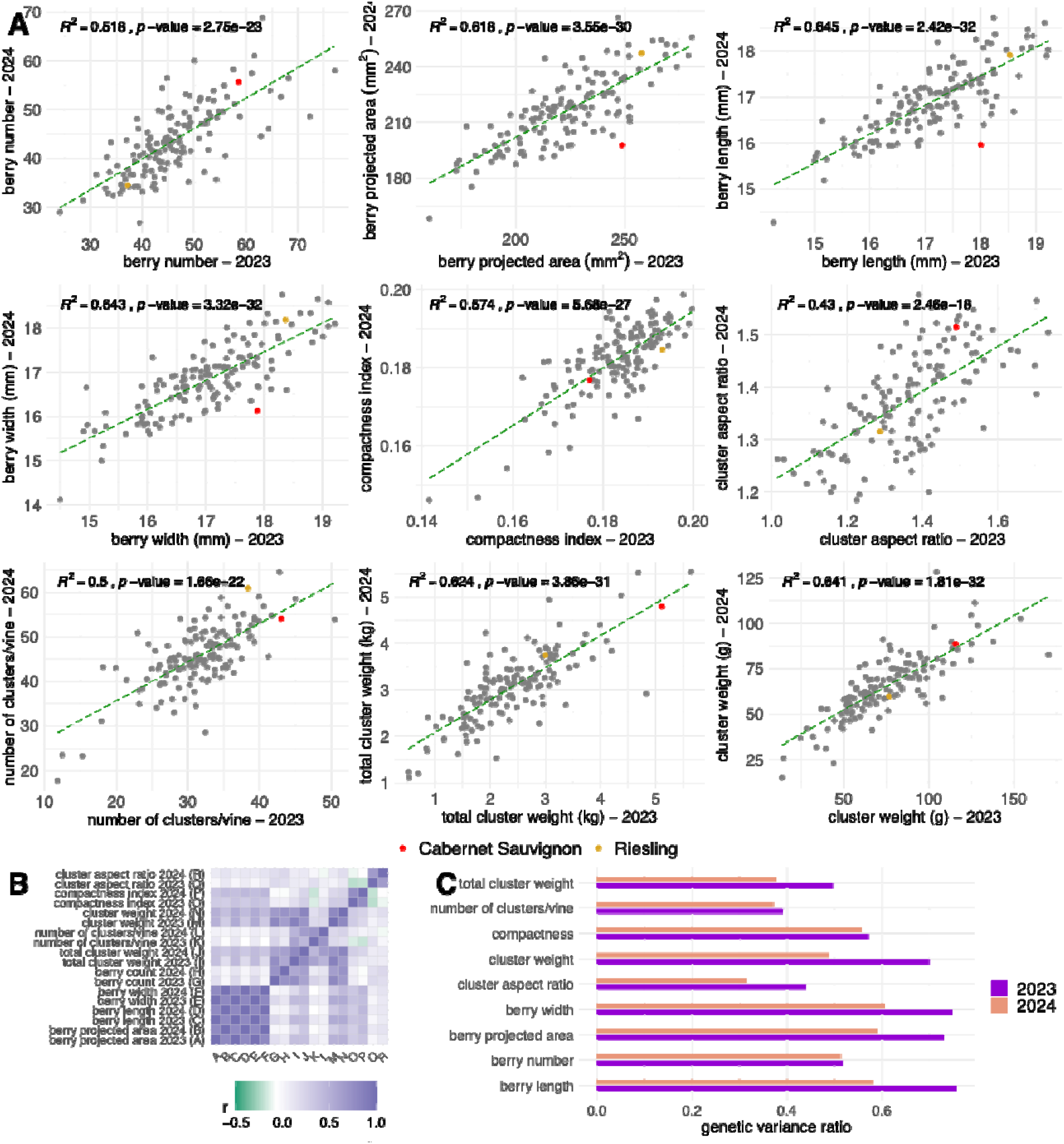
Phenotypic variability, trait correlations, and heritability. **(A)** Comparison of the 2023 and 2024 seasons for yield-related and digital traits based on BLUPs. The dashed green line represents the linear regression between the two seasons, while red and orange points correspond to the parental cultivars, Cabernet Sauvignon and Riesling, respectively. **(B)** Correlations between BLUPs for all traits were analyzed separately for each season. **(C)** Proportion of genetic variance relative to the total variance.

The number of clusters per vine increased from a mean of 31.58 ± 5.8 (SD) in 2023 to 45.62 ± 7.14 in 2024. Riesling and Cabernet Sauvignon followed this trend, increasing from 38.36 to 60.92, and from 42.98 to 54.09, respectively. These increases are likely attributable to differences in pruning and canopy management between years. In contrast, average cluster weight decreased slightly, from 74.94 g ± 25.98 in 2023 to 65.05 g ± 16.93 in 2024. Cabernet Sauvignon consistently produced heavier clusters than Riesling, with weights of 115.67 g versus 76.67 g in 2023, and 88.42 g versus 59.58 g in 2024. Total cluster weight per vine increased overall from 2.42 kg ± 0.86 in 2023 to 3.05 kg ± 0.77 in 2024. However, the parental cultivars exhibited different trends: Riesling increased from 3.00 to 3.73 kg, while Cabernet Sauvignon declined slightly from 5.11 to 4.79 kg, again likely reflecting differences in canopy management across seasons.

Among the image-derived traits, berry number per cluster averaged 46.02 ± 8.78 in 2023 and 43.65 ± 7.49 in 2024. Riesling produced fewer berries (37.2 in 2023, 34.37 in 2024), while Cabernet Sauvignon had higher counts (58.56 and 55.66, respectively). Berry projected area showed a slight decline from 222.41 mm^2^ ± 24.53 in 2023 to 215.99 mm^2^ ± 18.9 in 2024. Riesling maintained consistent berry sizes (∼248 mm^2^), while Cabernet Sauvignon showed a marked decrease from 257.75 mm^2^ in 2023 to 197.60 mm^2^ in 2024. Similar patterns were observed for berry length and width. Berry length averaged 17.08 mm ± 1.00 in 2023 and 16.89 mm ± 0.77 in 2024; berry width decreased from 17.02 mm ± 0.99 to 16.84 mm ± 0.79. Riesling remained relatively stable for these traits, while Cabernet Sauvignon showed mild reductions. Cluster compactness remained stable across years, with mean values of 0.18 ± 0.009 in both seasons. However, the parental lines were distinct: Riesling consistently exhibited greater compactness (0.19) than Cabernet Sauvignon (0.17), aligning with known phenotypic differences between these cultivars.

### Trait correlations and implications for cluster architecture

As expected, berry size-related traits were highly correlated (Figure 2B). For instance, berry projected area and berry length showed a very strong correlation (Pearson’s r = 0.99) in both years. Similarly, berry projected area was highly correlated with berry width (Pearson’s r > 0.98) in both years. Berry count exhibited a strong correlation with cluster weight (Pearson’s r = 0.81 in 2023 and 0.72 in 2024) and total cluster weight (Pearson’s r = 0.76 in 2023 and 0.62 in 2024). Additionally, the number of clusters per vine was positively correlated with total cluster weight (Pearson’s r is 0.60 in 2023 and 0.65 in 2024). Cluster compactness showed low to moderate correlations with berry area (Pearson’s r = 0.30 in 2023 and 0.41 in 2024), cluster weight (Pearson’s r = 0.30 in 2023 and 0.25 in 2024), total cluster weight (Pearson’s r = 0.16 in 2023 and 0.23 in 2024), berry length (Pearson’s r = 0.36 in 2023 and 0.44 in 2024), and berry width (r = 0.37 in 2023 and 0.45 in 2024). A negative correlation was observed between cluster aspect ratio and the compactness index in 2023 (Pearson’s r = -0.22), suggesting that their aspect ratio decreases as clusters become more compact, meaning they become wider and less elongated. Conversely, when clusters are more elongated, with a higher aspect ratio, berries are more loosely packed, reducing compactness. Furthermore, the compactness index for both years was negatively correlated with the number of clusters per vine (Pearson’s r = -0.15 in 2023 and -0.19 in 2024). This might suggest that when the number of clusters per vine decreases, individual clusters receive more resources, such as water, nutrients, and carbohydrates, leading to an increased number of berries per cluster and tighter packing, which results in a higher compactness index.

### Heritability of yield and digital traits

Heritability estimates varied across traits and years (Figure 2C). Berry number had a moderate heritability of 0.52 in 2023 and 0.51 in 2024. The berry’s projected area had higher heritability (0.73 in 2023, dropping to 0.59 in 2024), reflecting greater environmental influence in the second year. Berry length and width showed similar patterns, with heritability decreasing from 0.76 to 0.58 for length and from 0.75 to 0.61 for width. Cluster compactness showed moderate heritability (0.57 in 2023, 0.56 in 2024), while the number of clusters per vine had lower estimates (0.39 in 2023, 0.37 in 2024). Cluster weight showed a decline in heritability from 0.7 in 2023 to 0.49 in 2024, and total cluster weight followed a similar trend (0.5 in 2023, 0.38 in 2024), indicating increased environmental influence in 2024.

### QTL mapping for cluster compactness and yield components

Two genetic maps were constructed using a pseudo-test cross linkage mapping strategy [12]. The Cabernet Sauvignon parental map comprised 3,030 markers distributed across 982 bins (unique genetic positions), while the Riesling parental map contained 3,360 markers across 889 bins. Overall, physical and genetic distances showed strong collinearity, indicating a well-structured mapping framework (Figure 3).

**Figure 3.**
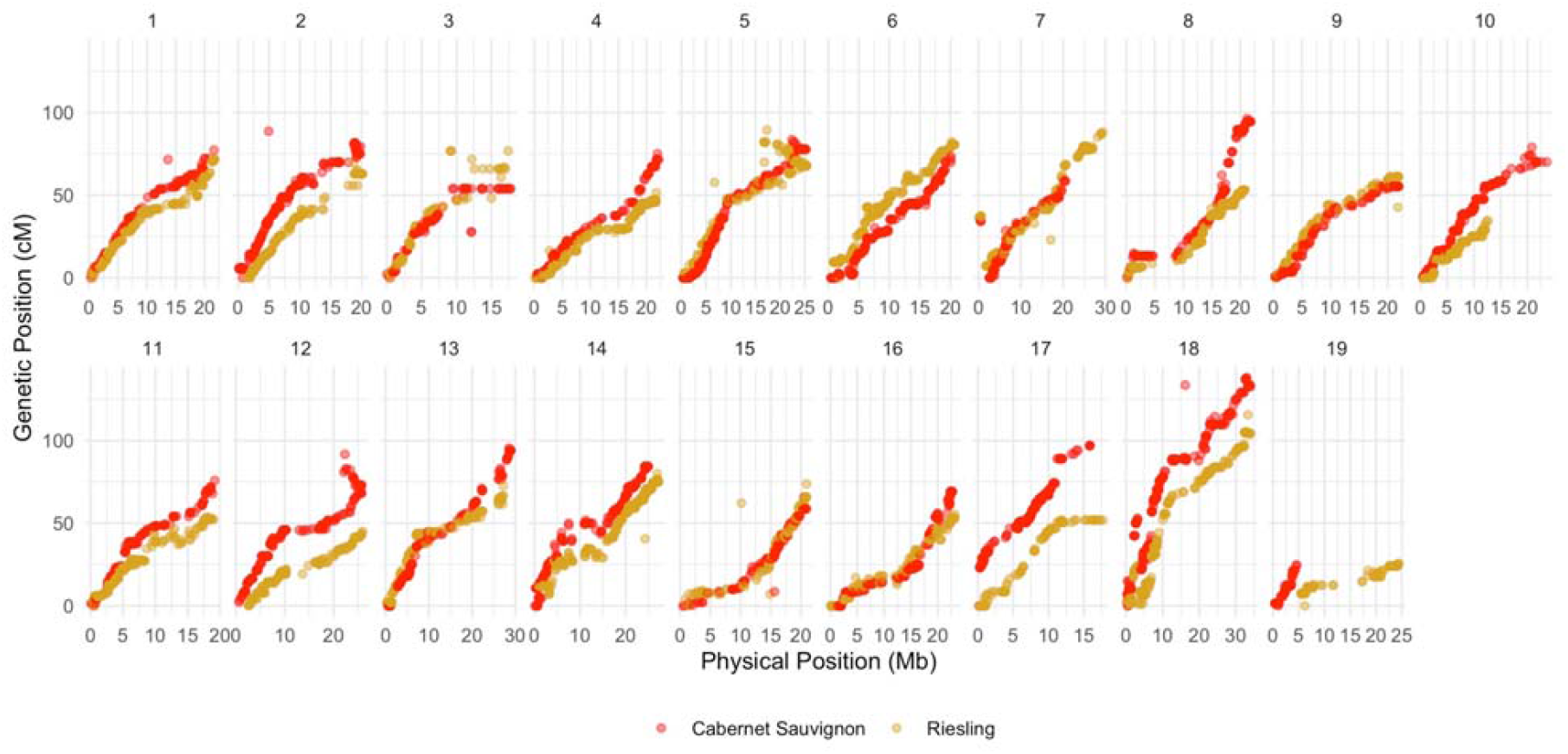
Relationship between genetic and physical positions for each chromosome in the parental maps.

QTL mapping identified nine consistent QTLs in both growing seasons (Figure 4). Eight QTLs were found on the Cabernet Sauvignon parental map with markers segregating as AB×AA or lm×ll, while one QTL was detected on the Riesling parental map with markers segregating as AA×AB or nn×np. These QTLs were associated with berry area, compactness, berry count, length, and width. Specifically, QTLs for berry length, width, and area were consistently detected on chromosomes 6 and 17 in Cabernet Sauvignon, with marker effects ranging from 7.6% to 17.8%. The co-localization of QTLs for these traits is likely due to the high correlation between these traits. A QTL for compactness was identified on chromosome 1 with marker effects of 10.3% in 2023 and 12.3% in 2024. Two QTLs on chromosomes 5 and 7 were found in Cabernet Sauvignon and Riesling, respectively, with marker effects between 9.2% and 22.1%.

**Figure 4.**
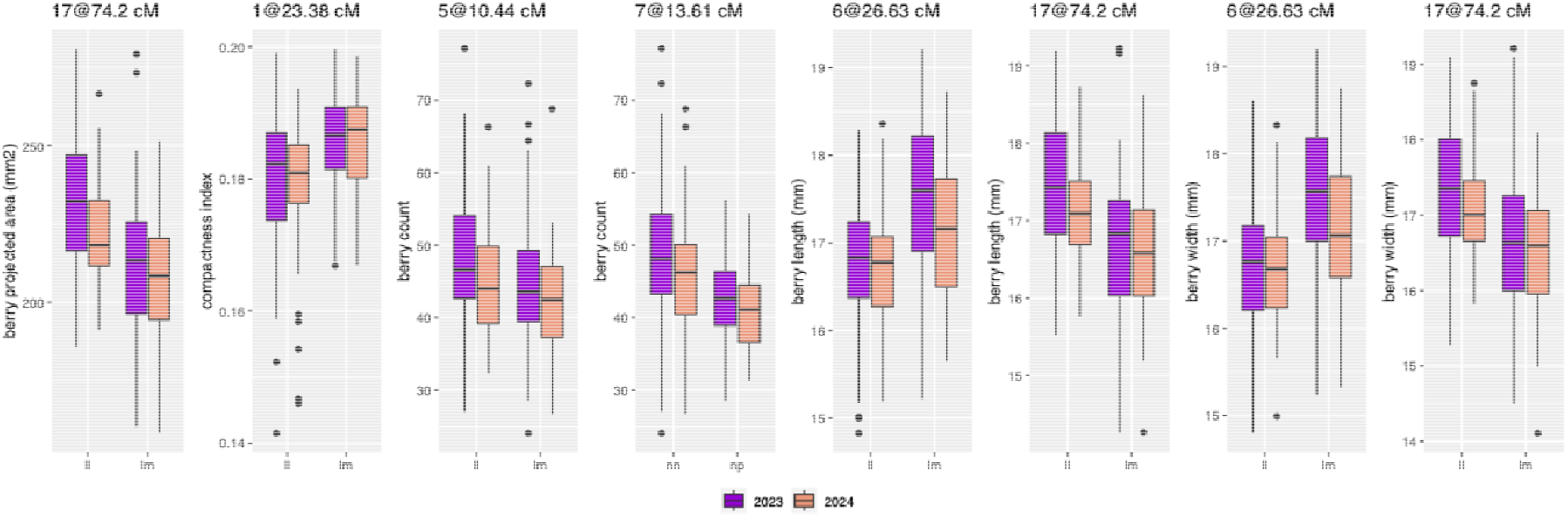
Marker effects for QTLs identified in two growing seasons. QTL mapping was conducted independently for both parents using a backcross model (pseudo-testcross QTL mapping). Therefore, markers are either (i) lm × ll, where Cabernet Sauvignon is heterozygous and Riesling is homozygous, or (ii) nn × np, where Cabernet Sauvignon is homozygous, and Riesling is heterozygous.

An additional 29 QTLs were identified using data from a single year, with marker effects of up to 22.9% (for number of clusters in the Riesling genetic map). The effect sizes for all QTLs, including those detected in one or both years, are summarized in Table 1.

**Table 1.** QTLs for yield-related traits and cluster compactness identified in a Riesling × Cabernet Sauvignon population using a pseudo-test cross linkage mapping strategy. QTLs detected in both years, based on the 1.5 LOD interval, are displayed in the top part of the table. Physical positions are based on the Cabernet Sauvignon cl.08 v1.1 haplotype 1 genome.

| Year | Parent | Trait | Chr | Genetic position (cM) | Physical position (bp) | Marker variance (%) | Model variance (%) | 1.5 LOD interval (cM) |
| --- | --- | --- | --- | --- | --- | --- | --- | --- |
| *Consistent QTL discovered in two years* |  |  |  |  |  |  |  |  |
| 2023 | Cab Sauv | berry area | 6 | 26.63 | 8,170,801 | 16.0 | 31.6 | 22.1 |
| 2024 |  |  |  | 21.58 | 6,208,069 | 13.2 | 64.1 | 11.9 |
| 2023 | Cab Sauv | berry area | 17 | 74.2 | 10,728,284 | 17.8 | 31.6 | 19.1 |
| 2024 |  |  |  | 62.69 | 8,241,035 | 15.9 | 64.1 | 18.4 |
| 2023 | Cab Sauv | compactness | 1 | 23.38 | 4,848,398 | 10.3 | 10.3 | 35.8 |
| 2024 |  |  |  | 21.47 | 4,241,270 | 12.3 | 12.3 | 31.5 |
| 2023 | Cab Sauv | berry count | 5 | 10.44 | 4,649,500 | 17.3 | 22.2 | 9.4 |
| 2024 |  |  |  | 10.08 | 4,379,291 | 19.3 | 23.4 | 9.4 |
| 2023 | Riesling | berry count | 7 | 13.61 | 3,144,729 | 22.1 | 38.0 | 10.7 |
| 2024 |  |  |  | 14.33 | 4,617,742 | 9.2 | 34.8 | 26.5 |
| 2023 | Cab Sauv | berry length | 6 | 26.63 | 8,170,801 | 15.2 | 32.1 | 22.1 |
| 2024 |  |  |  | 21.58 | 6,208,069 | 9.9 | 30.4 | 32.9 |
| 2023 | Cab Sauv | berry length | 17 | 74.2 | 10,728,284 | 19.1 | 32.1 | 18.8 |
| 2024 |  |  |  | 74.2 | 10,728,284 | 15.1 | 30.4 | 36.4 |
| 2023 | Cab Sauv | berry width | 6 | 26.63 | 8,170,801 | 14.9 | 29.7 | 21.1 |
| 2024 |  |  |  | 23.87 | 6,969,197 | 7.6 | 29.0 | 33.9 |
| 2023 | Cab Sauv | berry width | 17 | 74.2 | 10,728,284 | 16.8 | 29.7 | 20.2 |
| 2024 |  |  |  | 74.2 | 10,728,284 | 15.7 | 29.0 | 36.4 |
| Single year QTLs |  |  |  |  |  |  |  |  |
| 2024 | Cab Sauv | berry area | 1 | 30.05 | 5,783,345 | 5.6 | 64.1 | 46.5 |
| 2024 | Cab Sauv | berry area | 5 | 58.33 | 16,726,371 | 10.7 | 64.1 | 5.0 |
| 2024 | Cab Sauv | berry area | 8 | 76.26 | 18,524,489 | 5.2 | 64.1 | 44.4 |
| 2024 | Cab Sauv | berry area | 10 | 66.05 | 18,712,836 | 5.1 | 64.1 | 31.4 |
| 2024 | Cab Sauv | berry area | 11 | 49.46 | 11,515,400 | 14.2 | 64.1 | 13.1 |
| 2024 | Cab Sauv | berry area | 12 | 53 | 20,509,714 | 9.9 | 64.1 | 31.5 |
| 2024 | Cab Sauv | berry area | 14 | 22.82 | 2,731,011 | 6.8 | 64.1 | 12.6 |
| 2024 | Riesling | aspect ratio | 3 | 40.94 | 6,732,209 | 12.0 | 12.0 | 41.7 |
| 2023 | Cab Sauv | cluster weight | 5 | 2.88 | 2,842,257 | 10.0 | 21.2 | 13.7 |
| 2023 | Riesling | cluster weight | 7 | 12.18 | 3,144,650 | 16.5 | 38.5 | 10.7 |
| 2023 | Riesling | cluster weight | 13 | 56.51 | 21,438,556 | 14.2 | 38.5 | 25.0 |
| 2023 | Riesling | cluster weight | 14 | 62.83 | 22,578,129 | 6.1 | 38.5 | 40.8 |
| 2024 | Riesling | cluster weight | 14 | 10.61 | 3,135,853 | 11.6 | 11.6 | 74.3 |
| 2023 | Cab Sauv | cluster weight | 17 | 48.3 | 5,387,153 | 8.9 | 21.2 | 89.0 |
| 2024 | Cab Sauv | cluster weight | 17 | 74.2 | 10,728,284 | 11.6 | 11.6 | 55.8 |
| 2023 | Riesling | cluster weight | 18 | 7.81 | 4,773,569 | 7.0 | 38.5 | 67.7 |
| 2023 | Riesling | compactness | 11 | 23.01 | 5,263,713 | 12.6 | 12.6 | 45.2 |
| 2023 | Cab Sauv | berry count | 4 | 75.09 | 21,951,164 | 18.2 | 22.2 | 11.5 |
| 2024 | Cab Sauv | berry count | 4 | 67.17 | 21,001,269 | 20.1 | 23.4 | 21.6 |
| 2024 | Riesling | berry count | 6 | 60.05 | 14,808,244 | 7.3 | 34.8 | 76.6 |
| 2023 | Riesling | berry count | 13 | 55.44 | 20,915,884 | 17.7 | 38.0 | 14.3 |
| 2024 | Riesling | berry count | 14 | 10.61 | 3,135,853 | 16.2 | 34.8 | 21.8 |
| 2023 | Riesling | berry count | 18 | 11.72 | 6,123,609 | 10.6 | 38.0 | 35.7 |
| 2024 | Cab Sauv | berry length | 18 | 66.72 | 8,376,291 | 8.7 | 30.4 | 36.5 |
| 2023 | Riesling | number of clusters | 5 | 25.25 | 5,685,740 | 14.9 | 23.5 | 17.7 |
| 2023 | Riesling | number of clusters | 17 | 51.56 | 11,986,380 | 22.9 | 23.5 | 7.2 |
| 2023 | Riesling | total cluster weight | 7 | 9.32 | 3,001,260 | 10.4 | 22.3 | 17.9 |
| 2023 | Riesling | total cluster weight | 13 | 57.23 | 22,415,752 | 12.6 | 22.3 | 16.5 |
| 2024 | Cab Sauv | berry width | 18 | 65.84 | 8,317,021 | 8.9 | 29.0 | 36.5 |

## Discussion

This study offers novel insights into the genetic determinants of grape cluster architecture and yield components through QTL mapping in a bi-parental population derived from Riesling × Cabernet Sauvignon. Leveraging image-based phenotyping powered by the Segment Anything Model [13], or SAM, we acquired high-resolution morphological data for over 7,000 cluster images, leading to the identification of multiple QTLs associated with berry size and cluster compactness. Notably, our findings reveal stable genetic loci across multiple years, underscoring their potential applications in breeding programs aimed at enhancing grape quality and disease resistance.

Compact grape clusters have been widely associated with increased susceptibility to fungal diseases, particularly *Botrytis* bunch rot, due to restricted airflow, increased humidity retention, and elevated temperatures within the cluster [14]. Moreover, dense clusters inhibit the development of the protective waxy cuticle, a key barrier against pathogen invasion [15-16]. This compromised cuticle weakens the plant’s natural defense mechanisms, rendering berries more vulnerable to rot-fungal infections and pest damage [6]. Additionally, excessive berry compactness elevates the likelihood of physical damage, including berry cracking and juice leakage [15], creating an optimal environment for fungal spores to germinate and propagate, thereby exacerbating disease pressure [14]. Reduced sunlight exposure in compact clusters further limits UV-induced skin thickening and wax deposition, structural adaptations that enhance natural defense mechanisms against fungal infections [17-18]. Furthermore, heterogeneous ripening in compact clusters adversely affects sugar accumulation [19-20], anthocyanin biosynthesis [21-22], and tannin content [23-24], leading to inconsistent fruit maturity and diminished wine quality [25], ultimately resulting in economic losses for growers. To mitigate the adverse effects of compact clusters, growers commonly apply gibberellic acid GA to loosen clusters, especially in the table grape industry [26-27]. However, integrating genetic solutions through breeding presents a more sustainable and cost-effective approach.

A key contribution of this work is the application of foundational models like SAM to process images and generate high-quality masks for all berries within a cluster. Traditional methods for assessing grape cluster architecture include visual scoring, manual measurements, and conventional computer vision techniques, each with significant limitations. Visual scoring is subjective and prone to evaluator bias, manual measurements are labor-intensive and impractical for large-scale studies, and conventional computer vision models require extensive training and often fail to generalize across different grape varieties and imaging conditions. Cluster compactness is a critical trait that lacks easy and scalable measurement methods, yet it significantly influences grape quality, disease susceptibility, and yield. Several limitations commonly encountered when phenotyping cluster architecture and compactness can be effectively addressed with the cluster analysis pipeline presented in this study. For example, Underhill et al. (2020) highlighted how variation in peduncle length across genotypes affects the positioning of clusters when clipped for imaging [3]. Short peduncles can cause the tops of clusters to extend outside the camera’s field of view, leading to incomplete data capture. By using SAM, each object is segmented separately and then filtered based on size or shape, ensuring that only grape berries are retained for analysis. Since rachis and clippers have distinct morphometric features, they can be easily discarded, allowing clusters to be suspended centrally within the image frame to ensure that the entire cluster remains within the field of view. This is especially useful for highly shouldered clusters with short peduncles. Moreover, berry segmentation with SAM is equally effective regardless of berry color, addressing a common challenge in traditional threshold-based methods, which often struggle to separate objects of interest from the background. Unlike conventional methods, SAM, a zero-shot model, requires no task-specific training and is generalizable enough to perform effectively on grapevine clusters. This study demonstrates SAM’s high accuracy in identifying individual berries in two-dimensional cluster images, achieving a strong correlation with human-identified berries (Pearson’s r = 0.96). Using this model, we processed thousands of cluster images, generating nearly 330,000 berry masks, each linked to spatial coordinates within their clusters. This capability enables comprehensive, high-throughput analysis of cluster architecture, eliminating the need for time-consuming manual annotation while ensuring high reproducibility across different imaging conditions [2]. By leveraging SAM’s automated and scalable segmentation capabilities, we were able to accurately phenotype grape clusters and identify a moderate-effect QTL for cluster compactness across multiple harvest seasons in a pure *Vitis vinifera* population. This highlights the potential of foundation models in plant phenotyping and their ability to streamline genetic studies of complex traits.

These cultivars differ markedly across a range of biological and agronomic traits, including their geographical origins, historical patterns of cultivation, phenology, growth habits, fruit chemistry, and enological characteristics. Most relevant to this study, however, are their contrasting cluster architectures—Riesling typically forms smaller, more compact clusters, while Cabernet Sauvignon produces larger, looser ones. With a strong consumer preference for pure *V. vinifera* cultivars and wines, identifying trait donors within *V. vinife*ra is crucial for breeding strategies. Unlike breeding programs that rely on non-vinifera genetic material, which often necessitate multiple rounds of backcrossing to eliminate undesirable traits, the use of pure *V. vinifera* donors allows for more efficient gene introgression. While the individual marker effects identified in this study may not be strong enough to justify marker-assisted selection as a standalone strategy, our findings underscore the tremendous genetic variability for cluster architecture that can be unlocked through a single cross. This highlights the potential for recurrent selection, coupled with novel tools such as genomic prediction and AI-enabled computer vision, as a promising breeding approach to improve grape cluster structure and adaptability.

While this study provides valuable insights into the genetic architecture of grape cluster architecture and yield components, it is important to acknowledge some limitations such as the mapping population size of 138 individuals, which although sufficient for detecting moderate-effect QTLs, limits the resolution of QTL intervals and statistical power for identifying small-effect loci. Consequently, some QTL regions span relatively large genomic intervals, making it challenging to pinpoint individual candidate genes for each trait with confidence. Further studies incorporating larger populations and complimentary approaches such as fine mapping and transcriptomics could help narrow these intervals and refine gene discovery. Nonetheless, the stable QTLs detected across years in this study serve as promising targets for further validation and breeding efforts.

## Material and methods

### Plant material

The study utilized a genetic population of 138 full-sibling progeny derived from a cross between *Vitis vinifera* cv. Riesling and Cabernet Sauvignon. This cross was initially made by M.A. Walker and C.P. Meredith in 1994, and the progeny were initially planted at the Wolfskill Experimental Orchards in Winters, CA. In 2017, the plants were propagated, grafted onto 3309C, and transplanted to the UC Davis Viticulture and Enology Department Experimental Station in Oakville, Napa County, CA, USA (38°25′45.4′′N; 122°24′36.4′′W). This same population and planting have been used to understand the genetic basis of microbiome recruitment in grapevine [28]. The experimental design consisted of three randomized blocks, each containing three vines per genotype, leading to a total of nine replicates per genotype and 1,242 vines. Within each block, the three replicate plants of a genotype were placed consecutively in a row, with their positions randomized across the three blocks. The vineyard was established with 1.8 m × 2.4 m vine and row spacing, with rows oriented in a northwest-southeast direction. The vines were trained to a modified vertical shoot positioning system and pruned to two-bud spurs during winter dormancy. Drip irrigation was applied using two pressure-compensating emitters per plant. Cabernet Sauvignon vines were planted as a buffer surrounding the trial to isolate the experimental population from the vineyard edges.

### Traditional phenotyping

Grape clusters were harvested during two growing seasons, 2023 and 2024. In 2023, harvest took place on October 4; in 2024, it was conducted on September 20. All clusters from the middle vine of the three consecutive vines per genotype were collected for each genotype. The total number of clusters per vine was recorded, and the clusters were weighed to determine the total cluster weight per vine and average cluster weight. Additionally, five clusters were randomly selected from each middle vine for imaging, as described in the following section. Harvest was completed in a single day for each season to maintain consistency. Ripening time (i.e., technological maturity) varied across the population but was not recorded. Fruit was harvested when Cabernet Sauvignon, one of the parental varieties, reached approximately 24.5-25.5 °Brix.

### Digital phenotyping

To assess grape cluster architecture, imaging and image processing were performed following a previously developed methodology [2]. For each vine, five clusters were randomly selected and photographed using a Canon EOS 70D camera equipped with a 24-mm prime lens under standardized lighting conditions. All images were captured at a fixed distance, and a reference object was included in each frame to allow for size normalization and correction of any variations in camera-to-cluster distance. Camera settings were kept constant (aperture f/5, exposure time 1/500 s) to ensure image consistency. Based on previous findings, we captured two images per cluster, one from the front (0°) and one from the side (90°), as this approach has been shown to reliably estimate berry count and overall cluster structure [2]. Before segmentation, a region of interest (ROI) was automatically defined to isolate the grape cluster from the background. This step ensures that the segmentation algorithm processes only the relevant parts of the image. In this context, segmentation refers to the process of identifying and delineating each individual berry within the cluster image. This is a critical step that enables downstream analysis of cluster architecture based on detailed berry-level data.

For segmentation, we used the Segment Anything Model (SAM), a state-of-the-art computer vision tool that allows automated object identification. Within the defined ROI, a grid of 32 × 32 evenly spaced prompts guided the model to identify potential objects. The SAM framework, equipped with a ViT-H (Vision Transformer – Huge) encoder, generated precise segmentation masks that delineated individual berries within each image.

Post-processing steps were applied to refine the initial predictions. Segmented objects were filtered based on geometric properties such as area, perimeter, aspect ratio, and intersection-over-union (IoU) thresholds to remove incorrectly labeled structures. To further improve accuracy, elliptical Fourier descriptors were computed for each segmented shape, followed by principal component analysis (PCA) to distinguish true berries from non-berry elements like rachis fragments or background artifacts. This resulted in a clean and accurate set of berry masks suitable for phenotypic analysis. Final berry-level measurements, including berry count, size, shape, and spatial distribution, were extracted and analyzed using the R package Momocs [29]. All image processing and analysis steps were implemented in Python 3.11.

### Genotyping and linkage mapping

The mapping population, including both progeny and parents, was genotyped using genotyping-by-sequencing (GBS) following the method of Hyma et al. (2015) [30]. DNA was extracted from young leaves using the DNeasy 96-well DNA extraction kit (Qiagen, Valencia, CA, USA). Library preparation involved digestion with ApeKI, followed by sequencing on a HiSeq2000 platform (Illumina Inc., San Diego, CA, USA) at the Institute of Biotechnology, Genomics Facility (Cornell University, Ithaca, NY). Raw sequencing reads were processed using the TASSEL 5 GBS v2 pipeline with default parameters [31]. Single nucleotide polymorphisms (SNPs) were identified by aligning sequence reads to the Cabernet Sauvignon cl.08 v1.1 Hap1 [32] genome assembly using BWA [33-34]. The TASSEL 5 standalone program was then used to filter SNPs based on coverage depth (minimum read depth >6) and minor allele frequency (>0.1). SNPs and genotypes with more than 10% missing data were removed, and SNPs within 64 base pairs were merged.

Two genetic maps, one for Cabernet Sauvignon and another for Riesling, were constructed using a pseudo-test cross-linkage mapping strategy [12] with the R package BatchMap [35]. QTL mapping was performed using the R/qtl function stepwiseqtl, which applies forward and backward selection to identify a multiple QTL model [36]. Model selection was based on a penalized LOD score, with separate penalties for main effects and interactions. Penalties were calculated using 1,000 iterations for each trait.

## Statistical analysis

For traditional yield-related traits (number of clusters per vine, total cluster weight per vine, and average cluster weight), we used a mixed model with block as a fixed effect and genotype as a random effect. For the digital traits, including berry number, berry projected area, length, width, and cluster compactness, we first averaged the values across the five imaged clusters per genotype per block, then applied a similar mixed model with block and genotype as fixed and random effects, respectively. Best linear unbiased predictors (BLUPs) were estimated for each trait and used for QTL mapping. All statistical models were implemented using the R package lme4 [37].

## Acknowledgments

The authors would like to thank Guillermo Garcia-Zamora, Dan Ng, Mikayla Bailey, Gonzalo Ruiz, and Jovita Lopez for their support.

## Author information

### Authors and Affiliations

^1^Department of Viticulture and Enology, University of California Davis, CA 95616, USA

## Contributions

S.S, J.M, and L.D.G planned, designed the experiments, and wrote the manuscript. S.S, J.M, H.B, Y.L, V.N, A.G, and L.D.G collected the data and prelimimary analysis. E.T-L and L.D.G performed image analysis. S.S, J.M, and L.D.G analyzed genotypic data and QTL mapping. S.S, J.M, E.T-L, J. L, H.B, Y.L, V.N, A.G, D.C and L.D.G reviewed the manuscript.

## References

1. Tello, J., Aguirrezábal, R., Hernáiz, S., Larreina, B., Montemayor, M. I., Vaquero, E., & Ibáñez, J. (2015). Multicultivar and multivariate study of the natural variation for grapevine bunch compactness. Australian Journal of Grape and Wine Research, 21(2), 277–289. 10.1111/ajgw.12121

2. Torres-Lomas, E., Lado-Bega, J., Garcia-Zamora, G., & Diaz-Garcia, L. (2024). Segment Anything for comprehensive analysis of grapevine cluster architecture and berry properties. Plant Phenomics, 6, 0202. DOI:10.34133/plantphenomics.0202

3. Underhill, A., Hirsch, C., & Clark, M. (2020). Image-based phenotyping identifies quantitative trait loci for cluster compactness in grape. Journal of the American Society for Horticultural Science, 145(6), 363–373. 10.21273/JASHS04932-20

4. Vail, M. E., Wolpert, J. A., Gubler, W. D., & Rademacher, M. R. (1998). Effect of cluster tightness on Botrytis bunch rot in six Chardonnay clones. Plant Disease, 82(1), 107–109. 10.1094/PDIS.1998.82.1.107

5. Richter, R., Gabriel, D., Rist, F., Töpfer, R., & Zyprian, E. (2019). Identification of co-located QTLs and genomic regions affecting grapevine cluster architecture. Theoretical and Applied Genetics, 132, 1159–1177. 10.1007/s00122-018-3269-1

6. Herzog, K., Wind, R., & Töpfer, R. (2015). Impedance of the grape berry cuticle as a novel phenotypic trait to estimate resistance to Botrytis cinerea. Sensors, 15(6), 12498–12512. 10.3390/s150612498

7. Tello, J., & Ibáñez, J. (2018). What do we know about grapevine bunch compactness? A state-of-the-art review. Australian Journal of Grape and Wine Research, 24(1), 6–23. 10.1111/ajgw.12310

8. Zyprian, E., Richter, R., Rossmann, S., Theres, K., & Töpfer, R. (2018, July). Molecular analysis of bunch architecture in grapevine. In XII International Conference on Grapevine Breeding and Genetics 1248 (pp. 327–330). 10.17660/ActaHortic.2019.1248.47

9. Azevedo, C. F., Ferrão, L. F. V., Benevenuto, J., de Resende, M. D. V., Nascimento, M., Nascimento, A. C. C., & Munoz, P. R. (2024). Using visual scores for genomic prediction of complex traits in breeding programs. Theoretical and Applied Genetics, 137(1), 9. 10.1007/s00122-023-04512-w

10. Cubero, S., Diago, M. P., Blasco, J., Tardáguila, J., Prats-Montalbán, J. M., Ibanez, J., & Aleixos, N. (2015). A new method for assessment of bunch compactness using automated image analysis. Australian Journal of Grape and Wine Research, 21(1), 101–109. 10.1111/ajgw.12118

11. Correa, J., Mamani, M., Munoz-Espinoza, C., Laborie, D., Muñoz, C., Pinto, M., & Hinrichsen, P. (2014). Heritability and identification of QTLs and underlying candidate genes associated with the architecture of the grapevine cluster (Vitis vinifera L.). Theoretical and Applied Genetics, 127, 1143–1162. 10.1007/s00122-014-2286-y

12. Grattapaglia, D., & Sederoff, R. (1994). Genetic linkage maps of Eucalyptus grandis and Eucalyptus urophylla using a pseudo-testcross: mapping strategy and RAPD markers. Genetics, 137(4), 1121.10.1093/genetics/137.4.1121

13. Kirillov, A., Mintun, E., Ravi, N., Mao, H., Rolland, C., Gustafson, L., & Girshick, R. (2023). Segment anything. In Proceedings of the IEEE/CVF international conference on computer vision (pp. 4015–4026).

14. Hed, B., Ngugi, H. K., & Travis, J. W. (2009). Relationship between cluster compactness and bunch rot in Vignoles grapes. Plant Disease, 93(11), 1195–1201. 10.1094/PDIS-93-11-1195

15. Marois, J. J., Nelson, J. K., Morrison, J. C., Lile, L. S., & Bledsoe, A. M. (1986). The influence of berry contact within grape clusters on the development of Botrytis cinerea and epicuticular wax. American Journal of Enology and Viticulture, 37(4), 293-296. 10.5344/ajev.1986.37.4.293

16. Gabler, F. M., Smilanick, J. L., Mansour, M., Ramming, D. W., & Mackey, B. E. (2003). Correlations of morphological, anatomical, and chemical features of grape berries with resistance to Botrytis cinerea. Phytopathology, 93(10), 1263–1273. 10.1094/PHYTO.2003.93.10.1263

17. Percival, D. C., Sullivan, J. A., & Fisher, K. H. (1993). Effect of cluster exposure, berry contact and cultivar on cuticular membrane formation and occurrence of bunch rot(Botrytis cinerea PERS.: FR.) with 3 Vitis vinifera L. cultivars. Vitis, 32(2), 87–97.

18. Martínez-Lüscher, J., Torres, N., Hilbert, G., Richard, T., Sánchez-Díaz, M., Delrot, S., & Gomès, E. (2014). Ultraviolet-B radiation modifies the quantitative and qualitative profile of flavonoids and amino acids in grape berries. Phytochemistry, 102, 106–114. 10.1016/j.phytochem.2014.03.014

19. Pieri, P., Zott, K., Gomès, E., & Hilbert, G. (2016). Nested effects of berry half, berry and bunch microclimate on biochemical composition in grape. Oeno One, 50(3), 23–41. 10.20870/oeno-one.2016.50.3.52

20. Sweetman, C., Sadras, V. O., Hancock, R. D., Soole, K. L., & Ford, C. (2014). Metabolic effects of elevated temperature on organic acid degradation in ripening Vitis vinifera fruit. Journal of Experimental Botany, 65(20), 5975–5988. 10.1093/jxb/eru343

21. Sadras, V. O., & Moran, M. A. (2012). Elevated temperature decouples anthocyanins and sugars in berries of Shiraz and Cabernet Franc. Australian Journal of Grape and Wine Research, 18(2), 115–122. 10.1111/j.1755-0238.2012.00180.x

22. Rienth, M., Torregrosa, L., Luchaire, N., Chatbanyong, R., Lecourieux, D., Kelly, M. T., & Romieu, C. (2014). Day and night heat stress trigger different transcriptomic responses in green and ripening grapevine (Vitis vinifera) fruit. BMC Plant Biology, 14, 1–18. 10.1186/1471-2229-14-108

23. Cohen, S. D., Tarara, J. M., & Kennedy, J. A. (2008). Assessing the impact of temperature on grape phenolic metabolism. Analytica Chimica Acta, 621(1), 57–67. 10.1016/j.aca.2007.11.029

24. Koyama, K., Ikeda, H., Poudel, P. R., & Goto-Yamamoto, N. (2012). Light quality affects flavonoid biosynthesis in young berries of Cabernet Sauvignon grape. Phytochemistry, 78, 54–64. 10.1016/j.phytochem.2012.02.026

25. Armstrong, C. E. J., Ristic, R., Boss, P. K., Pagay, V., & Jeffery, D. W. (2021). Effect of grape heterogeneity on wine chemical composition and sensory attributes for Vitis vinifera cv. Cabernet Sauvignon. Australian Journal of Grape and Wine Research, 27(2), 206–218. 10.1111/ajgw.12469

26. Domingos, S., Nobrega, H., Raposo, A., Cardoso, V., Soares, I., Ramalho, J. C., & Goulao, L. F. (2016). Light management and gibberellic acid spraying as thinning methods in seedless table grapes (Vitis vinifera L.): Cultivar responses and effects on the fruit quality. Scientia Horticulturae, 201, 68–77. 10.1016/j.scienta.2016.01.034

27. Wegher, M., Faralli, M., & Bertamini, M. (2022). Cluster-zone leaf removal and GA3 application at early flowering reduce bunch compactness and yield per vine in Vitis vinifera cv. Pinot Gris. Horticulturae, 8(1), 81. 10.3390/horticulturae8010081

28. Floerl, L., Lin, J., Griggs, R. G., Massonnet, M., Cochetel, N., Figueroa-Balderas, R., Cantu, D., & Bokulich, N. A. (2025). The genetic basis of microbiome recruitment in grapevine and its association with fermentative and pathogenic taxa. bioRxiv. 10.1101/2025.03.31.646331

29. Bonhomme, V., Picq, S., Gaucherel, C., & Claude, J. (2014). Momocs: outline analysis using R. Journal of Statistical Software, 56, 1–24. 10.18637/jss.v056.i13

30. Hyma, K. E., Barba, P., Wang, M., Londo, J. P., Acharya, C. B., Mitchell, S. E., & Cadle-Davidson, L. (2015). Heterozygous mapping strategy (HetMappS) for high resolution genotyping-by-sequencing markers: a case study in grapevine. PLoS One, 10(8), e0134880. 10.1371/journal.pone.0134880

31. Glaubitz, J. C., Casstevens, T. M., Lu, F., Harriman, J., Elshire, R. J., Sun, Q., & Buckler, E. S. (2014). TASSEL-GBS: a high capacity genotyping by sequencing analysis pipeline. PloS one, 9(2), e90346. 10.1371/journal.pone.0090346

32. Massonnet, M., Cochetel, N., Minio, A., Vondras, A. M., Lin, J., Muyle, A., & Cantu, D. (2020). The genetic basis of sex determination in grapes. Nature communications, 11(1), 2902. 10.1038/s41467-020-16700-z

33. Li, H., & Durbin, R. (2009). Fast and accurate short read alignment with Burrows–Wheeler transform. Bioinformatics, 25(14), 1754–1760. 10.1093/bioinformatics/btp324

34. Chin, C. S., Peluso, P., Sedlazeck, F. J., Nattestad, M., Concepcion, G. T., Clum, A., & Schatz, M. C. (2016). Phased diploid genome assembly with single-molecule real-time sequencing. Nature Methods, 13(12), 1050–1054. 10.1038/nmeth.4035

35. Schiffthaler, B., Bernhardsson, C., Ingvarsson, P. K., & Street, N. R. (2017). BatchMap: a parallel implementation of the OneMap R package for fast computation of F1 linkage maps in outcrossing species. PloS One, 12(12), e0189256. 10.1371/journal.pone.0189256

36. Broman, K. W., Wu, H., Sen, Ś., & Churchill, G. A. (2003). R/qtl: QTL mapping in experimental crosses. Bioinformatics, 19(7), 889–890. 10.1093/bioinformatics/btg112

37. Bates, D., Mächler, M., Bolker, B., & Walker, S. (2015). Fitting linear mixed-effects models using lme4. Journal of Statistical Software, 67, 1–48. 10.18637/jss.v067.i01

